# New RoxS sRNA targets identified in *B. subtilis* by pulsed SILAC

**DOI:** 10.1101/2023.02.17.528916

**Authors:** Delphine Allouche, Gergana Kostova, Marion Hamon, Christophe H. Marchand, Mathias Caron, Sihem Belhocine, Ninon Christol, Violette Charteau, Ciarán Condon, Sylvain Durand

**Author notes:** These authors contributed equally to this work (Delphine Allouche, Gergana Kostova).

## Abstract

Non-coding RNAs (sRNA) play a key role in controlling gene expression in bacteria, typically by base-pairing with ribosome binding sites to block translation. The modification of ribosome traffic along the mRNA generally affects its stability. However, a few cases have been described in bacteria where sRNAs can affect translation without a major impact on mRNA stability. To identify new sRNA targets in *B. subtilis* potentially belonging to this class of mRNAs, we used pulsed-SILAC (stable isotope labeling by amino acids in cell culture) to label newly synthesized proteins after short expression of the RoxS sRNA, the best characterized sRNA in this bacterium. RoxS sRNA was previously shown to interfere with the expression of genes involved in central metabolism, permitting control of the NAD+/NADH ratio in *B. subtilis*. In this study, we confirmed most of the known targets of RoxS, showing the efficiency of the method. We further expanded the number of mRNA targets encoding enzymes of the TCA cycle and identified new targets primarily regulated at the translational level. One of these is YcsA, a tartrate dehydrogenase that uses NAD+ as co-factor, in excellent agreement with the proposed role of RoxS in Firmicutes.

**Importance:** Non-coding RNA (sRNA) play an important role in bacterial adaptation and virulence. The identification of the most complete set of targets for these regulatory RNAs is key to fully identify the perimeter of its function(s). Most sRNAs modify both the translation (directly) and mRNA stability (indirectly) of their targets. However, sRNAs can also influence the translation efficiency of the target primarily, with little or no impact on mRNA stability. The characterization of these targets is challenging. We describe here the application of the pulsed SILAC method to identify these targets and obtain the most complete list of targets for a defined sRNA.

## Introduction

The importance of *trans-*acting small regulatory RNAs (sRNA) in post-transcriptional regulation has been widely demonstrated in bacteria. The early characterization of sRNAs was mostly done in *Escherichia coli* and its pathogenic relatives. However, in the past decade, an increasing number of sRNAs have also been characterized in Gram-positive bacteria, such *Bacillus subtilis* and its virulent kin. These studies have highlighted common features and interesting differences. For example, most sRNAs in Gram-negative bacteria characterized thus far require either the Sm-like protein Hfq or the RNA chaperone ProQ to efficiently bind to their targets (review in (1)). However, ProQ is absent from Gram-positive bacteria and, although Hfq is present, this protein is not generally required for efficient base-pairing and regulation of most sRNAs characterized so far, with relatively few exceptions (2).

In most cases, sRNAs bind at or close to the ribosome binding site (RBS) and inhibit mRNA translation. This often indirectly results in mRNA destabilization, because ribosomes no longer protect the transcript from attack by ribonucleases (RNases). Small RNAs can also bind outside of the RBS and directly provoke mRNA degradation, by changing RNA structure or by directly recruiting RNases (3, 4). sRNAs can also positively regulate the expression of some targets by modifying the secondary structure of the 5’ untranslated region (5’-UTR) to release the RBS and increase translation efficiency, or change structure to impair RNase cleavages (reviewed in (5)).

Very few cases have been described in bacteria where sRNAs act at the translational level without affecting mRNA stability (6). A recent study in *S. aureus* identified two targets of the IsrR RNA, induced during iron starvation, that are primarily affected at the translational level (7). In all known cases, the sRNA base-pairs with the RBS.

The favored technique for global study of the effects of sRNAs on translation has been to compare proteomes in WT *versus* a strain deleted for the sRNA of interest. The main drawback of this approach is that it only provides a picture of the proteome at equilibrium, with numerous potential indirect effects. This issue has been partially circumvented by inducing sRNA expression for short periods of time (5 to 15 mins induction), allowing direct or indirect repressive effects on mRNA levels to be detected due to their short half-lives. However, the half-lives of proteins can be much longer than these induction times, making down-regulation of translation much more difficult to measure and potentially missed entirely in cases where the indirect effect on mRNA stability is minimal.

In this work, we have combined RNA sequencing (RNAseq) with pulsed SILAC (stable isotope labeling by amino acids in cell culture), first used in eukaryotes to identify miRNA targets (8), with a goal of identifying new sRNA targets in *B. subtilis* that potentially include some targets with primarily translational effects. To benchmark the approach, we chose to overexpress the RoxS sRNA, one of the best characterized sRNAs in Gram-positive bacteria, for a short period of time. RoxS is conserved in both *B. subtilis* and *S. aureus*, and uses three single-stranded C-rich regions (CRR1-3) to regulate the expression of genes involved in central metabolism. In *B. subtilis*, its main proposed function is to adjust temporary imbalances in the NAD+ to NADH ratio, by controlling key steps in fermentation pathways and the TCA cycle. Several direct targets of this sRNA have already been identified by the methods described above, allowing us to validate the performance of the pulsed SILAC experiment. In addition to identifying most of the known RoxS targets, we discovered several new mRNAs potentially regulated by this sRNA. Indeed, our data suggests that RoxS down-regulates the expression of all but one enzyme of the TCA cycle, a far greater impact than previously appreciated. As anticipated, we also identified some potential targets that were primarily regulated at the translational level, with only minor effects on mRNA stability. The potential repercussions of this type of regulation are discussed.

## Results

Pulsed SILAC consists in stable isotopic labeling by amino acids of newly synthetized proteins to allow their detection and quantification by mass spectrometry (MS). We performed experiments in a lysine auxotrophic strain to maximize the efficiency of lysine isotope incorporation into new proteins. The strain lacked the native version of the *roxS* gene and expressed RoxS under the control of an arabinose-inducible promoter from the *amyE* locus. The control strain carried the same arabinose promoter inserted at the *amyE* locus with the RoxS sRNA replaced by a transcriptional terminator to avoid polar effects on downstream genes. To ensure appropriate expression of mRNA targets, both strains were cultivated in controlled medium (MD) containing malate, a carbon source we have previously shown strongly induces RoxS expression (9). All 20 amino-acids in their light isotopic form were supplied to the medium, but lysine was limited to an empirically determined concentration that allowed it to be almost completely consumed by mid-exponential phase (OD_600_=0.6), leading to a growth arrest unless further supplemented with lysine isotopes (Supplementary Figure 1). At this point, arabinose and medium-heavy lysine were added to the control strain, and arabinose and heavy lysine added to the RoxS overexpressing strain, to simultaneously induce RoxS expression and label newly synthesized proteins. After 15 mins induction, bacteria were harvested to extract total RNA and soluble proteins. RNA was used for RNAseq analysis and equal amounts of extracted proteins were mixed and subjected to quantitative proteomic analysis. MS-based quantitative data (medium to light, heavy to light and medium to heavy ratios) determined at the peptide level for each lysine-containing peptide were further used to relatively quantify newly synthesized proteins in both conditions and determine those that were differently expressed upon RoxS induction.

The up- and down-regulated genes identified by RNAseq and by MS analysis after 15 min induction of RoxS are presented in Figure 1, Table 1 and 2. Only statistically significant alterations in expression (P-value ≤0.05) with a fold-change <0.7 and >1.4 for negatively and positively regulated targets respectively, were selected. This threshold includes all previously confirmed targets of RoxS in *B. subtilis*. As developed below, most of the new potential targets identified fit perfectly well with the known function of RoxS in the regulation of the NAD+/NADH ratio and its role in carbon metabolism (9–11).

**Figure 1:**
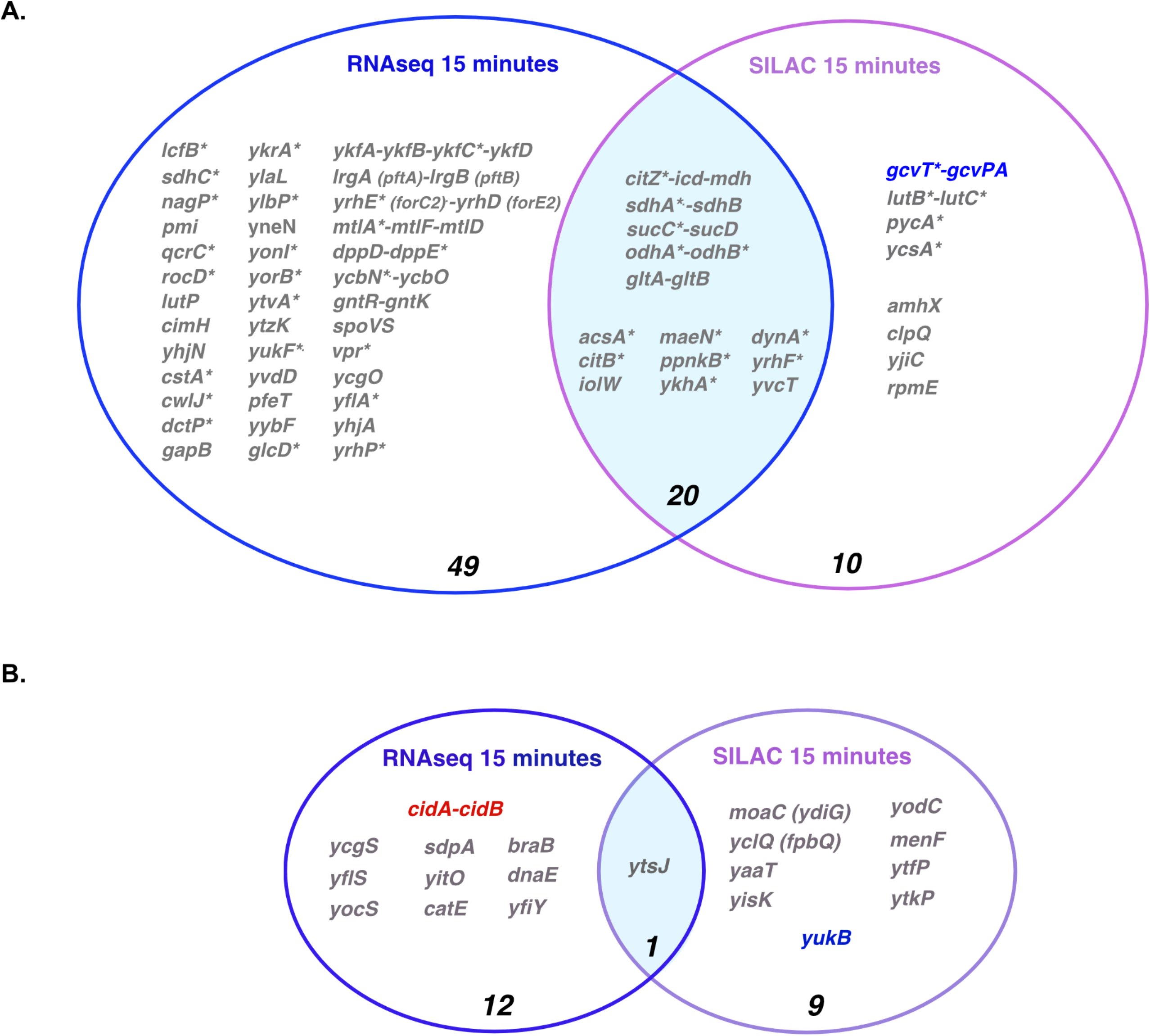
Venn diagrams of genes whose expression is altered at mRNA and protein levels after 15 minutes of RoxS overexpression. (**A**.) Downregulated genes. (**B**.) Upregulated genes. The number of genes identified by RNaseq, SILAC or both is indicated. (*) indicates genes with a predicted or demonstrated binding site for RoxS in the Shine-Dalgarno sequences. Candidates where no base-pairing could be predicted between the mRNA and RoxS by IntaRNA are shown in red (see Materials and Methods). mRNAs shown in blue were present in the RNaseq dataset, but the fold change was just below the threshold. These mRNAs thus potentially belong in the intersection of the Venn diagram.

**Table 1 and 2:**
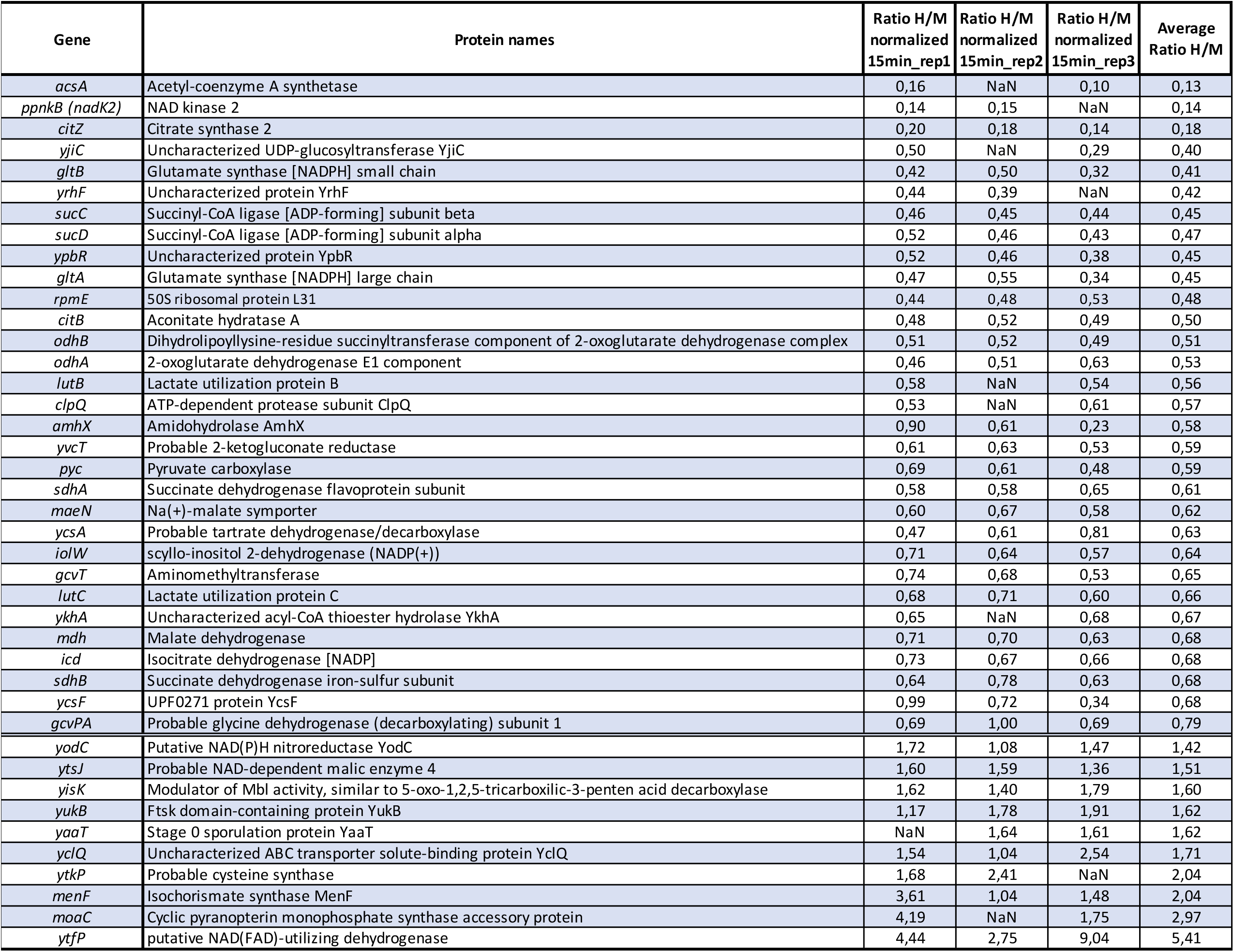
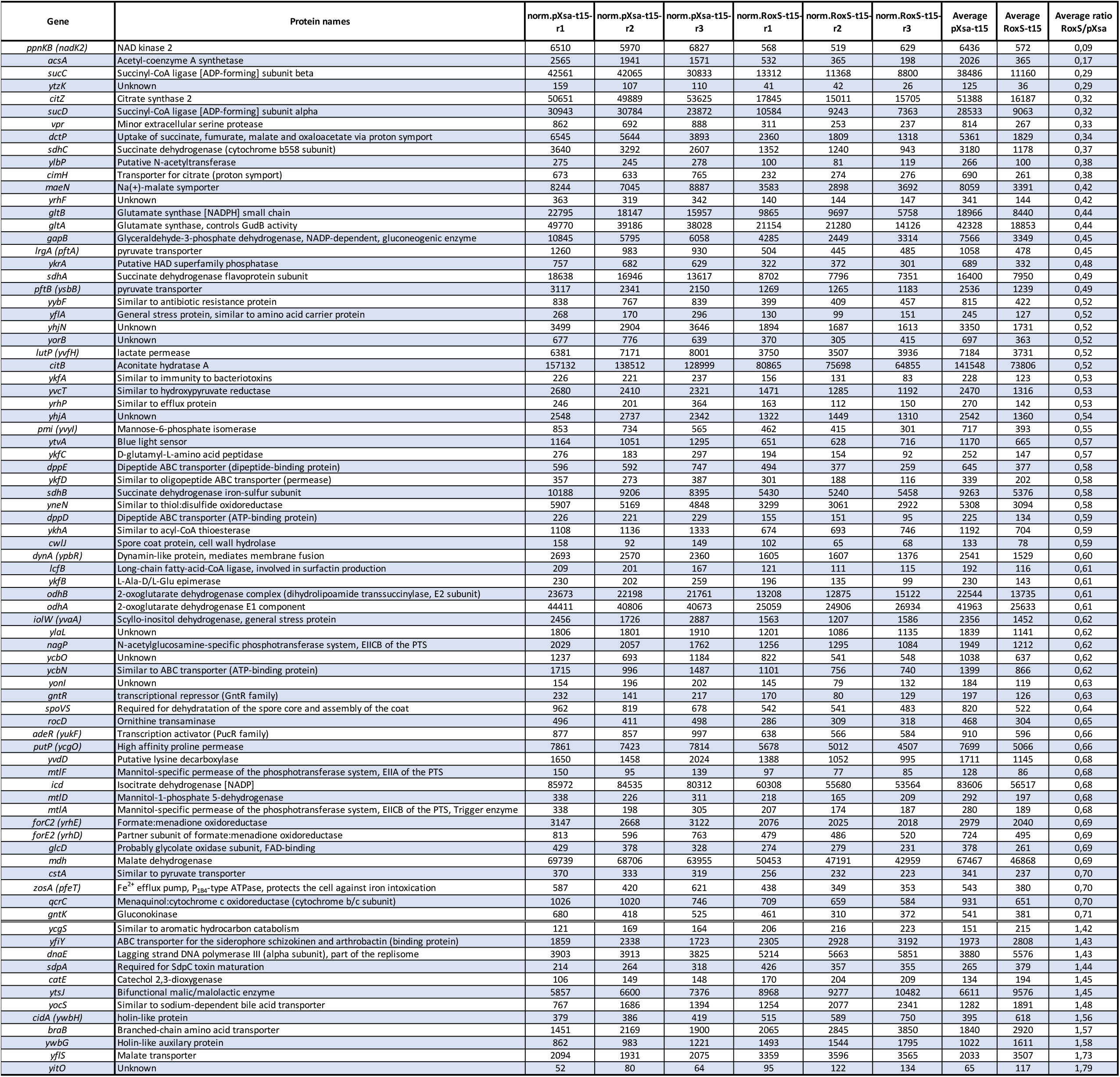
The up- and down-regulated genes identified by RNAseq (Table 2) and by MS analysis (Table 1) after 15 min induction of RoxS. Only statistically significant alterations in expression (P-value ≤0.05) with a fold-change <0.7 and ?1.4 for negatively and positively regulated targets respectively are presented.

### Genes whose expression is down-regulated by RoxS

In most cases described in literature, bacterial sRNAs that affect translation also indirectly alter mRNA stability. In agreement with this, we identified 21 potential targets (20 down-regulated and 1 up-regulated) that were found in both the RNAseq and the pulsed SILAC data sets (Figure 1). Among these candidates, were numerous previously identified targets of RoxS, such as *ppnkB* (an inorganic polyphosphate/ATP-NAD kinase), *ykhA* (a putative acyl-coenzyme A thioesterase), *acsA*, (an acetyl-CoA synthetase) and the gene encoding the TCA enzyme succinate dehydrogenase, *sucCD* (9–11). Interestingly, we identified 8 additional genes encoding enzymes of the TCA cycle, present on 4 different mRNAs (*citZ*-*icd-mdh, citB, odhAB, sdhAB*). All of these potential targets have at least one predicted base-pairing site for RoxS on the mRNA (Supplementary Figure 2). A Gene Ontology (GO) analysis of the data confirmed the over-representation of the TCA cycle genes (p-value of 1×10^−10^), compared to other cellular functions (Figure 2) (12). These results are in agreement with our previous data suggesting that RoxS regulates cell metabolism in order to limit NADH production (11). To confirm a representative example of these TCA cycle targets at the level of mRNA stability, we measured the half-life of the *citZ-icd-mdh* mRNA in a wild-type (WT) and *ΔroxS* mutant strain (Figure 3A). Two major species of the *citZ* mRNA were detected by Northern blot. The larger (∼4kb) band of corresponds to the tri-cistronic *citZ-icd-mdh* mRNA and the lower band (∼1kb) to the mono-cistronic *citZ* transcript. Two degradation products were also visible (D1 and D2), with D2 disappearing in the *ΔroxS* background, suggesting it is stabilized by the presence of RoxS (Figure 3A). The half-life of *citZ-icd-mdh* mRNA increased 3-fold in a Δrox*S* strain (3.3 min in a WT strain *vs* 9.6 min in a *ΔroxS* mutant) (Figure 3A). This result is in good agreement with the effect of RoxS measured by RNAseq (3-fold), but less than the effect detected by SILAC, where the level of the CitZ protein was reduced 5-fold upon over-expression of RoxS (Table 1), suggesting that translational control has a greater impact for this mRNA.

**Figure 2:**
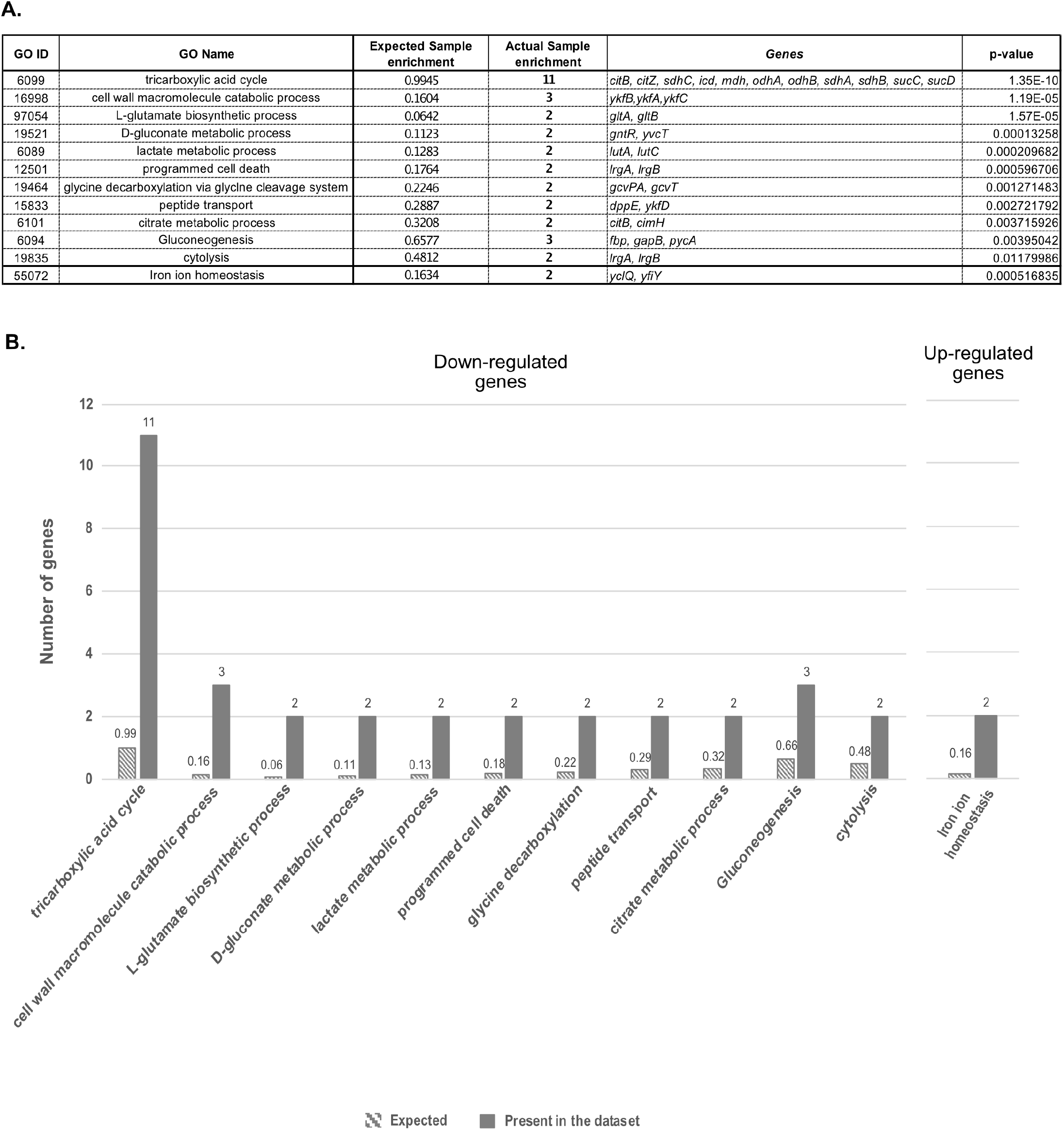
GO analysis of genes whose expression is altered at mRNA and protein levels after 15 minutes of RoxS overexpression. Analysis was performed using the Comparative GO web application (12). (**A**.) Table presenting the different biological functions over-represented in the dataset of potential RoxS targets with a p-value <0.05 (**B**.) Histogram representing the number of genes belonging to each biological function are over-represented in the down-regulated (left graph) and up-regulated (right graph) datasets.

**Figure 3:**
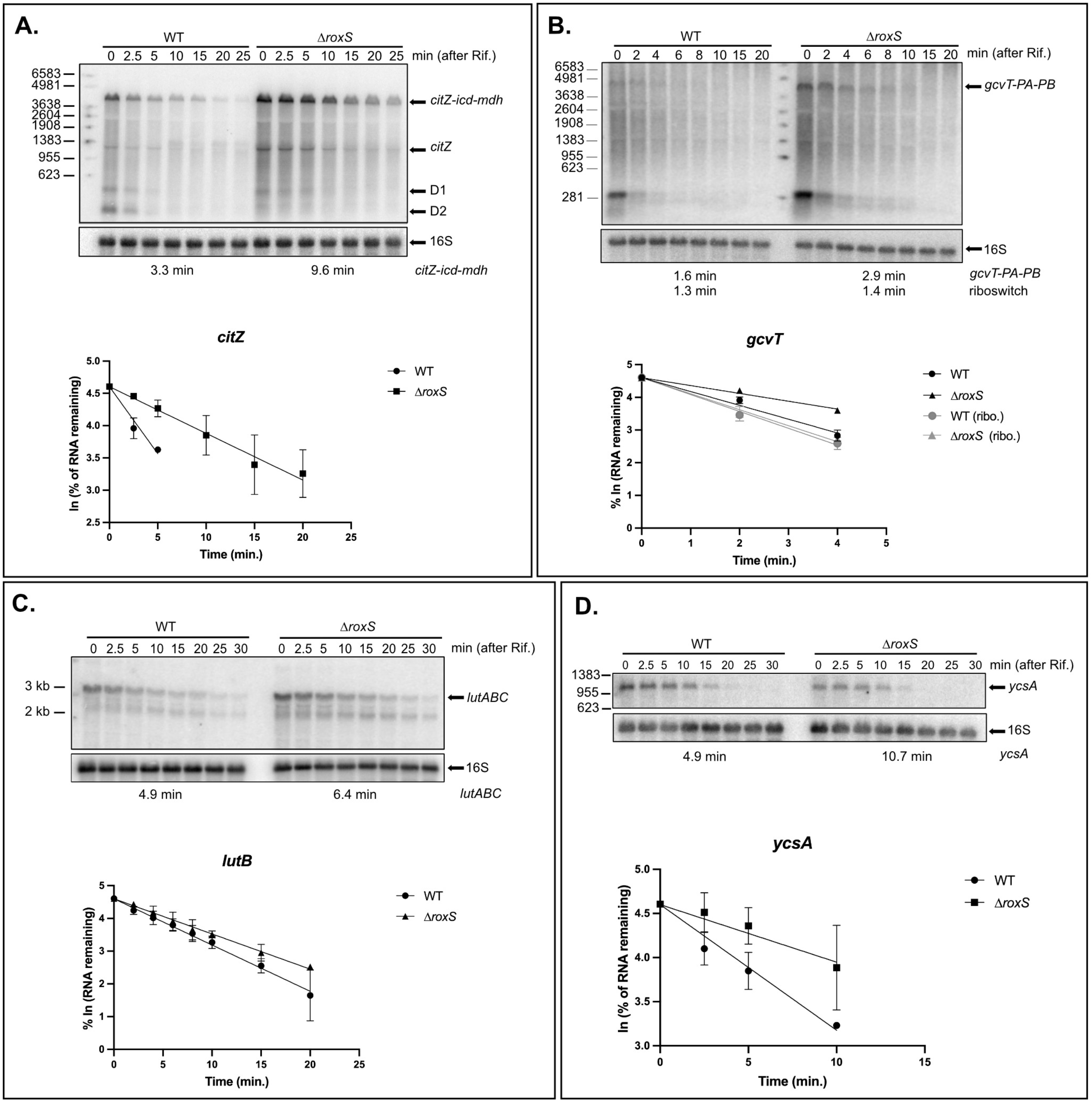
Half-life measurements of candidate RoxS target mRNAs. Northern blot of total RNA isolated from WT and *ΔroxS* mutant strain at different times after addition of rifampicin. Cells were grown in MD + malate (0.5%) and probed for (**A**.) *citZ*, (**B**.) *gcvT*, (**C**.) *lutB*, or in LB + malate (0.5%) and probed for (**D**.) *ycsA*. D1, D2: degradation products. Northern blots were re-probed for 16S rRNA as a loading control. Graphs and calculated from two independent experiments are shown beneath the autoradiographs.

The glycine decarboxylation pathway was also over-represented (p-value of 1.2×10^−6^) in the GO analysis of the potential target genes identified by pulsed SILAC (Figure 2). The level of the GcvT protein was reduced 1.5-fold after RoxS induction and the *gcvT-PA-PB* mRNA was reduced 1.3-fold, just below the threshold. The levels of second protein of the operon, GcvPA, were also reduced 1.3-fold. Several putative base-pairing sites for RoxS can be predicted on the *gcvT-PA-PB* mRNA, including in the SD regions of *gcvT* (supplementary Figure 2) and *gcvPB*, suggesting the operon is a *bona fide* target despite the weak effects. The stability of this mRNA increased 1.8-fold in the absence of RoxS (1.6 min in a WT strain *vs* 2.9 min in a *ΔroxS* strain; Figure 3B). Thus, this operon belongs to the category of targets where the effect on translation likely has a similar-magnitude indirect effect on mRNA stability, despite its position in the Venn diagram (Figure 1). Interestingly, the steady-state level of a ∼300-nt transcript corresponding to the glycine riboswitch was also increased in the *ΔroxS* strain. However, the half-life (∼1.3 min) was similar in both strains suggesting that RoxS may also have an indirect impact on the transcription of this operon.

Among the downregulated targets that were only identified by pulsed SILAC (and thus potentially regulated primarily at the translational level) were *lutB* and *lutC* of the *lutABC* operon, *ycsA* and *pycA*, all with prospective binding sites for RoxS in their SD regions (Supplementary Figure 3, Figure 4), and four genes (*amhX, clpQ, rpmE*, and *yjiC*), all with potential binding sites for RoxS in their ORFs (supplementary Figure 4). In the following paragraphs, we will describe the verification of the weak effect of RoxS on the half-lives of two of these potential targets, the *lutABC* and *ycsA* mRNAs, and additional experiments performed on *ycsA* to validate the effect on translation and the predicted RoxS binding site.

**Fig 4:**
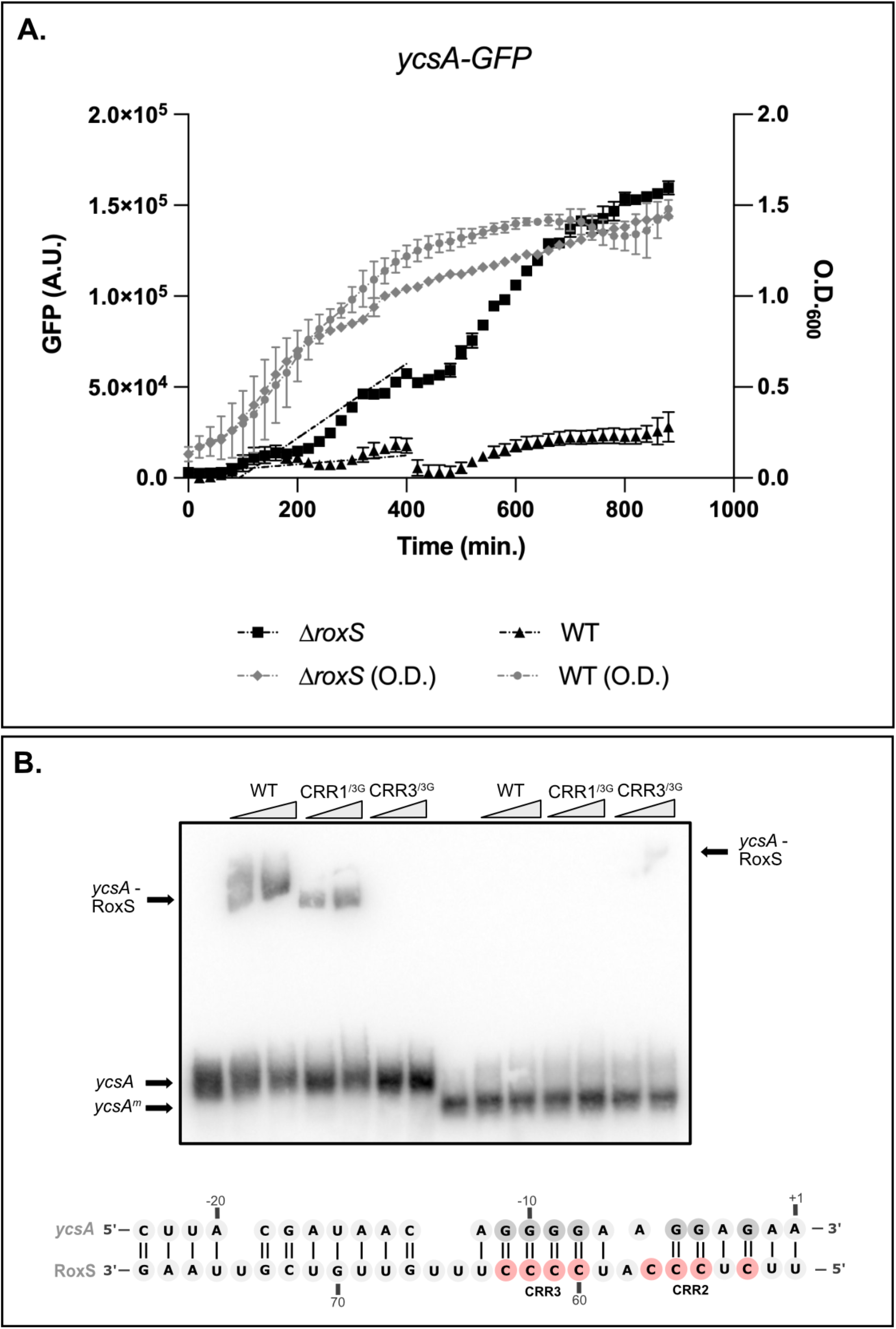
RoxS binds to the *ycsA* Shine-Dalgarno region to regulate translation. **A**. Measurement GFP fluorescence produced by the *ycsA*-GFP translational fusion during growth in LB + malate (0.5%), in the presence or absence of RoxS. **B**. EMSA experiment between the RoxS sRNA and the 5’ UTR of the *ycsA* mRNA. The binding of RoxS mutated in CRR1 and/or CRR3 (CRR1^/3G^, CRR3^/3G^) was also tested. Predicted base-pairing between the SD sequence of *ycsA* and C-rich region 2 and 3 of RoxS is shown below the autoradiogram.

The *lutABC* operon encodes lactate utilization enzymes, and interestingly is also regulated in *B. subtilis* by FsrA, an iron-induced sRNA with similar single-stranded C-rich motifs to RoxS involved in the binding of G-rich Shine-Dalgarno (SD) sequences (13). While FsrA was proposed to base-pair with the SD sequence of *lutA* (13), IntaRNA (14) predicts that both RoxS and FsrA could potentially bind close to the ribosome binding site of each of the three genes of this operon (Supplementary Figure 3). The levels of the LutB and LutC proteins were downregulated 1.8-fold and 1.5-fold respectively upon RoxS overexpression in the pulsed SILAC experiment. The effect of RoxS on mRNA stability was tested by measuring the impact of the RoxS deletion on the half-life of the *lutABC* mRNA by Northern blot (Figure 3C). In agreement with the RNAseq data, its half-life was relatively unaffected by the deletion of RoxS (4.9 min in a WT strain *vs* 6.4 min a *ΔroxS* strain; Figure 3B), suggesting the effect of RoxS occurs primarily at the translational level. The regulation of the *ycsA* gene, encoding a putative NAD-dependent tartrate dehydrogenase fits well with the proposed function of RoxS in managing NAD+/NADH ratios. The YcsA protein was reduced 1.6-fold upon RoxS induction in the pulsed SILAC experiment in MD medium. To obtain additional evidence for the translational regulation of *ycsA* by RoxS, we fused the *ycsA* ORF in frame to GFP and determined its rate of fluorescence production in the presence or absence of RoxS. This experiment was done in LB since the *ycsA* mRNA was shown to be much more expressed in these conditions from a previous study by Nicolas *et al*. (15) (Figure 4A). The rate of fluorescence accumulation during exponential phase increased more than 8-fold in a strain deleted for RoxS, confirming the translational regulation of *ycsA* expression by RoxS. We also determined the effect of the *ΔroxS* deletion on the stability of the *ycsA* mRNA in LB. Although the *ycsA* mRNA half-life increased 2.2-fold in a Δrox*S* strain compared to the WT strain (4.9 min in the WT strain *vs* 10.7 min in the *ΔroxS* mutant; Figure 3D), the translational effect on the fusion was much more pronounced that than the effect on the stability of this mRNA.

RoxS is predicted to interact with the ribosome binding site of *ycsA* mRNA primarily by base-pairing between four C residues of C-rich region CRR3 and four G-residues of the *ycsA* SD sequence (Figure 4B). To confirm this interaction, we performed an electromobility shift assay (EMSA) using an *in vitro*-transcribed fragment of *ycsA* containing the SD sequence and variants of RoxS where 3 residues of the C-rich regions 1 or 3 were mutated to G-residues (Figure 4B). The *ycsA* RNA formed a complex with RoxS and with the RoxS^CRR1^ mutant, but failed to interact with the RoxS^CRR3^ mutant, consistent with the prediction by IntaRNA. The introduction of the compensatory mutation in *ycsA* mRNA, where 3 G’s of the SD sequence were replaced by 3 C’s, abolished the interaction with WT RoxS, but partially restored the interaction with RoxS CRR3^3G^ mutant, supporting the interaction site prediction.

### Genes whose expression is up-regulated by RoxS

Only one potential positively regulated RoxS target, *ytsJ*, was identified by both RNAseq and pulsed SILAC (Figure 1). YtsJ is thought to be the primary malate dehydrogenase of four such enzymes identified in *B. subtilis* (16). In a previous study we showed that RoxS expression is induced by addition of malate to the growth medium and that RoxS positively regulates the expression of a second malate transporter (YflS) by protecting its mRNA from degradation by RNase J1 (9). Of note, the YflS protein was not detected by pulsed SILAC in this study, due to its localization in the membrane (see next section).

Previous studies have suggested that *ytsJ* is expressed from three different promoters, located in front of the upstream gene *dnaE* (*PdnaE*), in the middle of the *dnaE* ORF (*P2*) (15) or immediately upstream of *ytsJ* (*P1*) (Figure 5A) (16). We confirmed the 5’ ends of the *P2*-*ytsJ* and P1-*ytsJ* transcripts within the *dnaE* ORF by primer extension (supplementary Figure 5). These 5’ ends fit well with the promoter sequences previously proposed (15, 16). Interestingly, a binding site for RoxS was predicted by IntaRNA within the *dnaE* ORF, approximately 190 nts downstream of the *P2* promoter (Figure 5A). A second RoxS binding site within the YtsJ coding sequence (CDS) was also predicted, with a lower energy score.

**Figure 5:**
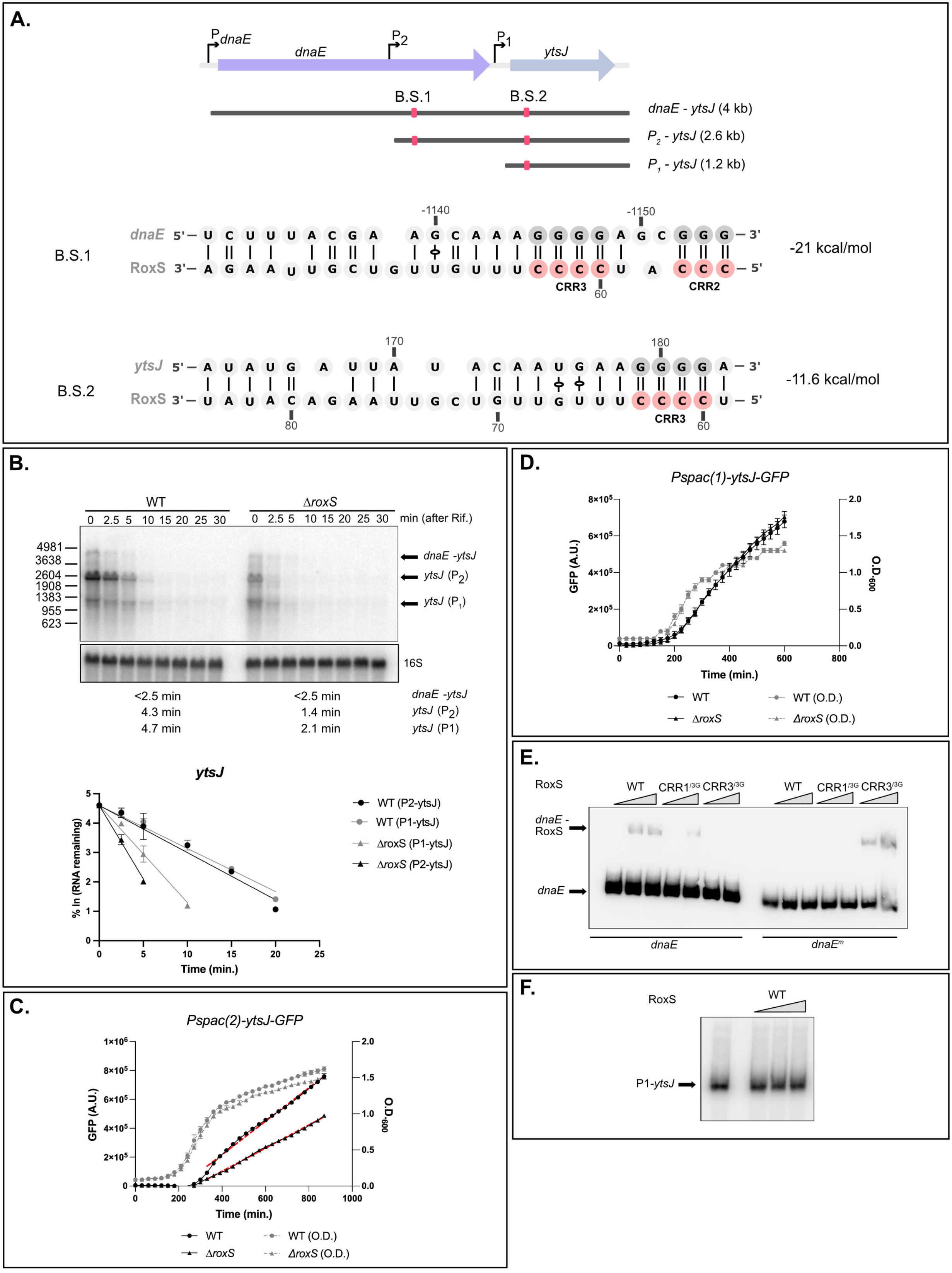
RoxS regulates expression of the P2*-ytsJ* transcript. **A**. Genomic environment of *ytsJ* gene. The three promoters mapped in this region PdnaE, P2 and P1 are shown as black arrows. Red squares indicates the locations of the predicted RoxS binding sites 1 (B.S.1) and 2 (B.S.2), with the interacting sequences shown. **B**. Northern blot of total RNA from WT and *ΔroxS* mutant strain isolated at different times after addition of rifampicin. Cells were grown in 2xTY + malate (0.5%) and probed for *ytsJ* mRNA. The blot was re-probed for 16S rRNA as a loading control. Graphs and calculated from two independent experiments are shown beneath the autoradiographs. (**C**.) and (**D**.) Fluorescence (black lines) produced by the Pspac2-*ytsJ-*GFP and Pspac1-*ytsJ-*GFP translational fusions, respectively, during growth (grey dashed lines) in LB + malate (0.5%) of WT and *ΔroxS* cells. (**E**.) EMSA experiments between WT and mutant variants of RoxS sRNA (CRR1^/3G^, CRR3^/3G^) and the P2*-ytsJ* mRNA transcript. (**F**.) EMSA experiments between the RoxS sRNA and P1-*ytsJ* mRNA.

We first measured the half-lives of the various *ytsJ* transcripts in a WT versus a ΔroxS strain grown in the same medium as the RNAseq experiment (MD + malate) by Northern blot using a riboprobe against the first half of the *ytsJ* coding sequence. Surprisingly, none of the three expected *ytsJ* primary transcripts, or an additional species of unknown origin, showed a measurable decrease in half-life (supplementary Figure 6), despite the 1.4-fold increase in mRNA levels found by RNAseq (Table 2). Since our previous studies had shown that RoxS regulation can depend on growth medium (9, 10), one possible explanation was greater RoxS expression under control of the arabinose-dependent promoter in the RNAseq experiment compared to that naturally occurring in MD + malate. (9, 10). Indeed, in rich medium (2xTY + malate), we were able to confirm the stabilizing effect of RoxS on *ytsJ* transcripts (Figure 5B). The half-life of the predominant *P2-ytsJ* (∼2.7 kb) transcript was reduced 2-fold in the *ΔroxS* strain compared to WT (5.3 *vs* <2.5 min). A similar effect was observed with the transcript corresponding in size to *P1-ytsJ* (5.6 *vs* <2.5 min), while no impact was seen on the bicistronic *dnaE-ytsJ* species (Figure 5B).

We next tested the effect of RoxS on the translation efficiency of *ytsJ* mRNA, by constructing two GFP fusions, expressed under the control of a Pspac promoter to remove complications of transcriptional effects from the native promoters. These fusions have a transcriptional start close to the P1 (*Pspac(1)-ytsJ-gfp*) or P2 promoter (*Pspac(2)-ytsJ-gfp)*. The experiments were done in LB + malate to reduce background fluorescence from the 2xYT medium. Consistent with the pulsed SILAC data where the YtsJ protein expression was induced 1.5-fold by RoxS, the rate of fluorescence accumulation of the P2-YtsJ-GFP fusion was 1.5-fold lower in the *ΔroxS* strain compared to WT, confirming that RoxS increases YtsJ expression (Figure 5C). However, we were unable to detect an impact of RoxS on the *P1-ytsJ-gfp* fusion (Figure 5D).

To determine whether RoxS is able to bind P2-*ytsJ* and P1-*ytsJ* transcripts, we tested the ability of RoxS to bind to RNA fragments containing the two predicted binding sites, by gel shift assay (Figure 5E). As expected, RoxS was able to bind to an RNA fragment within the *dnaE* ORF containing the binding site downstream of P2 and this interaction was abolished with the RoxS^CRR3^ mutant. A compensatory mutation in *dnaE* (*dnaE*^m^) restored base-pairing with the RoxS CRR3 mutant, confirming their ability to interact *via* the RoxS CRR3 region. We were unable to detect binding between RoxS and a fragment within the *ytsJ* ORF, containing the second potential binding site downstream of P1 (Figure 5F), consistent with the lack of regulation seen with the P1*-ytsJ-gfp* fusion. Together, these results suggest that RoxS directly regulates *ytsJ* expression on the P2 transcript, but not that from P1. The explanation for the effect of RoxS on the half-life of the *P1-ytsJ* transcript is unclear.

The position of the RoxS binding site far upstream (app. 1 kb) of the *ytsJ* start codon on the P2 transcript suggested that the primary effect might be on mRNA stability. To determine whether RoxS protects the P2-*ytsJ* mRNA against RNase J1 as observed previously for the upregulated *yflS* mRNA (9), or against RNase Y, one of the main endoribonucleases involved in mRNA degradation in *B. subtilis*, we analyzed the degradation profile of *ytsJ* mRNAs in *ΔrnjA and Δrny* mutant strains in presence or absence of RoxS (Figure 6A and B). These experiments were done in rich medium (2XTY) where the P2-*ytsJ* transcript is best detectable. The steady-state levels and stability of P2-*ytsJ* (and indeed all *ytsJ* transcripts) were no longer sensitive to RoxS in the *Δrny* mutant 13.9 min in both strains), suggesting that RoxS cannot stabilize *ytsJ* without prior action of RNase Y (Figure 6A and B). In the *ΔrnjA* mutant, up to seven different degradation intermediates were detected compared to the WT strain (D1-D7; Figure 6A). Two of these were impacted by the deletion of the *roxS* gene; D2 disappeared and was replaced by an intensified D3. This suggests that RoxS protects D2 from degradation by RNase J1. To clarify how RoxS might protect D2 from RNase J1, we performed a primer extension assay in the different RNase mutant strains to identify the 5’ ends of the degradation products (supplementary Figure 5). Since the signal detected from the native *ytsJ* gene was very low, we overexpressed a part of the 5’ UTR of the *P2-ytsJ* transcript to increase the intensity of the bands (supplementary Figure 5). The 5’ end corresponding to D2 was highly intensified in a *ΔrnjA* mutant strain and was absent in strains lacking either RoxS or RNase Y, suggesting that D2 is generated by RNase Y cleavage and protected from RNase J1 degradation by RoxS. Interestingly, this 5’ end is only few nucleotides (3-4) upstream of the RoxS binding site, a compatible location to block the processive activity of RNase J1 (17), similarly to what we previously observed for the *yflS* mRNA (18). The primer extension profile in the region of the second potential RoxS binding site in the *ytsJ* ORF was insensitive to the presence of RoxS in the different RNase mutant strains, reinforcing our conclusion that RoxS does not bind in the CDS of the *ytsJ* mRNA.

**Figure 6:**
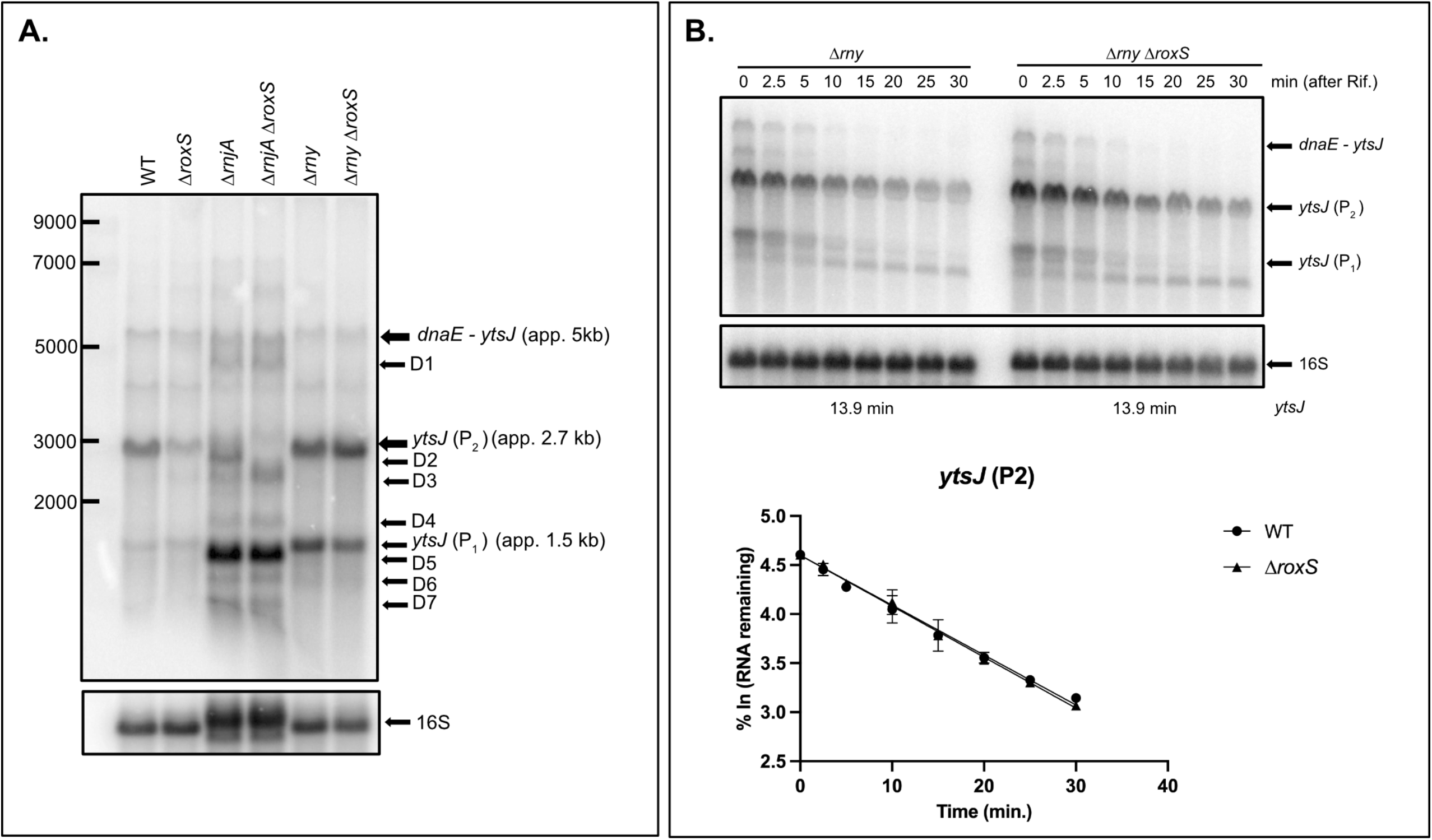
RoxS protects an RNase Y cleavage product of the P2-*ytsJ* transcript from degradation by RNase J. (A.) Northern blot showing various transcripts and degradation products of *ytsJ* detected in WT, *ΔroxS, ΔrnjA, ΔrnjA ΔroxS, Δrny* and *Δrny ΔroxS* strains. The Northern blot was re-probed for 16S rRNA as a loading control (note that a 16S processing intermediate accumulates in the rnjA background). (B). Northern blot of total RNA from *Δrny* and *Δrny ΔroxS* mutant strain isolated at different times after addition of rifampicin. Cells were grown in 2XTY + malate (0.5%) and probed for *ytsJ* mRNA. The blot was re-probed for 16S rRNA as a loading control. *ytsJ* (P1) and (P2) indicate *ytsJ* transcript from the P2 and P1 promoter, respectively. Graphs and calculated from two independent experiments are shown beneath the autoradiographs.

### Targets identified only by RNaseq

A number of potential targets were identified by RNAseq, but not by pulsed SILAC. All of these mRNAs are predicted to encode membrane proteins that would not have been present in the soluble protein fraction analyzed in the pulsed SILAC experiment. IntaRNA predicted RoxS binding site(s) in all cases but one (the *cidAB* operon), with energy scores of <-5 kcal/mol (see Materials and Methods). Although these interactions remain to be validated, it is interesting that almost 50% of the down-regulated targets identified by RNaseq, have a classical predicted binding site in their SD region. One up-regulated target, *yfiY* mRNA, which encodes a siderophore-binding subunit of an ABC transporter, has a predicted binding site for RoxS at its extreme 5’-end. The position of this binding site is similar to that described for *yflS* mRNA, where RoxS protects from the degradation by the 5’-3’ exoribonuclease J1 (9). These base-pairing predictions reinforce our belief that several of these membrane protein-encoding mRNAs are also potentially direct targets of RoxS (Figure 1).

## Discussion

This work shows that pulsed SILAC is an efficient alternative to identify direct effects of sRNAs on the translation of target genes, in particular down-regulatory effects that would normally be masked by protein stability. It is less cumbersome than ribosome-profiling experiments and has the major advantage of only focusing on newly synthesized proteins, without the background noise of translation occurring under equilibrium conditions. Using this technique, we not only identified most of the known targets of RoxS, but we also identified new potential targets for this sRNA in *B. subtilis*. We observed a clear enrichment of down-regulated genes involved in the TCA cycle, in agreement with the proposed function of RoxS to limit flux through this pathway to restrict NADH production in the presence of favorable carbon sources such as glucose or malate. We also identified the *ytsJ* mRNA as a new positively regulated target of RoxS. This gene encodes one of the four malate hydrogenases in *B. subtilis*, the only one that uses NADP+ instead of NAD+ as cofactor. The preferential up-regulation of an enzyme using NADP+ instead of NAD+ as cofactor fits well with the need for the cell to control NADH production. The enrichment of downregulated genes involved in metabolic pathways directly connected to the TCA cycle such as L-glutamate biosynthesis, citrate metabolism, glycine decarboxylation, and gluconeogenesis also is also consistent with the proposed metabolic role of RoxS (Figure 2).

This study also allowed us to identify *ycsA* mRNA as a direct target of RoxS, primarily regulated at its translational level. Indeed, we were able to measure a much greater impact of RoxS on translation (8-fold) than on the mRNA stability (2-fold). Of note, the weaker translational effect of RoxS measured by pulsed SILAC (1.6-fold) is probably explained by a lower translation efficiency of *ycsA* mRNA in MD medium. The fact that *ycsA* mRNA encodes a putative tartrate dehydrogenase that uses NAD+ as cofactor is also in good agreement with the proposed function of RoxS. A potential advantage of this type of regulation is the reversibility of control. Indeed, removal of the sRNA from its targets could allow new rounds of translation on negatively regulated targets or reducing the translation of positively regulated mRNAs but in keeping a basal level of expression. This impact on translation without affecting the pool of the mRNA could be a way to limit the need of new rounds of transcription and quickly adapt to changing environment. A parallel can be drawn to the situation in eukaryotes, where translationally repressed RNAs are stored in P-bodies to be recycled when needed (19).

In contrast to the *ycsA* mRNA, which has a classical binding site for RoxS in its SD region, several mRNAs that were regulated primarily at the translational level by RoxS, have at least one putative binding site predicted in their coding sequence (CDS) instead of the SD region, which is rather unusual (supplementary Figure 7). A few examples have been described in bacteria where the sRNA binds to the CDS well downstream (>5 codons) of the AUG. Some of these sRNAs impact mRNA stability with a likely indirect affect their translation (3, 20). In other cases, it has been shown that sRNA binding within the CDS can prevent formation of a secondary structure that stimulates translation initiation, lowering the translation efficiency of the target mRNA (21). The presence of sRNA binding sites deep within the CDS raise the question of how base-pairing occurs on an actively translated mRNA. Although we were able to confirm that RoxS has no impact on the mRNA stabilities of two mRNAs from this class of potential targets with putative RoxS binding sites within their ORFs: *yisK* (involved in the control of cell division) and *yjiC* mRNA (encoding a potential macrolide glycosyltransferase), we were unable to see an impact of RoxS on translation of these genes using a GFP-*yjiC* fusion or an antibody raised against YisK protein (data not shown). Furthermore, although the bicistronic *dnaE-ytsJ* and P2-*ytsJ* transcripts contain the RoxS binding site localized in *dnaE*, only the stability of the P2-*ytsJ* mRNA affected by RoxS (Figure 5), suggesting that translation of *dnaE* competes with RoxS regulation on the longer mRNA. We therefore cannot rule out that some mRNAs of this class may be off-targets due to the overexpression of RoxS. That said, some targets, such as *yukB*, probably merit further investigation since 9 potential binding site for RoxS are predicted within this CDS. One could imagine that transitory slow-downs in translation promoted by sRNA binding throughout the CDS could impact the folding of sub-domains of proteins without a major impact on the protein yield.

## Materials and Methods

### Strains, plasmids, oligonucleotides

Strains and plasmids used in this study are presented Table S1 and oligonucleotides in Table S2. The *lysA::erm* mutation was transferred by transformation of SSB1002 with chromosomal DNA from the strain BKE23380 (22). Plasmid pDG1662-PxsA (pl801) is a plasmid containing the arabinose inducible promoter of *xsa* gene, amplified using oligonucleotides CC2250 and CC2251. The PCR fragment was cloned between the BamHI and HindIII sites of pDG1662.

Plasmid pDG1662-PxsA-RoxS (pl805) was made by amplification of RoxS by PCR using oligos CC2285/CC2286 and the resulting fragment cloned between the SpeI and EcoRI restriction sites in pl801. The pDG148-*ycsA*-GFP (pl862) plasmid was made by PCR amplification of *ycsA* from chromosomal DNA using oligonucleotides CC2546/CC2547 and *gfp* from pHM2-5’hbs-GFP plasmid (pl696) using oligonucleotides CC2548/CC572. The two overlapping PCR fragments containing *ycsA* and *gfp* were assembled in new PCR reaction with oligo pair CC2546/CC572. The *ycsA*-GFP fragment was cloned in pHM2 between the EcoRI and BamHI restriction sites. Due to the low expression after insertion of this fusion at the *amyE* locus, this construction was reamplified with oligo pair CC2611/CC2612 to clone it in pDG148 between the EcoRI and PvuII restriction sites.

To overexpress the part of *dnaE* predicted to base-pair with RoxS, we inserted pHM2-Pspac^con^-*dnaE* at the *amyE* locus. This plasmid was constructed by amplification of a part of *dnaE* by PCR using oligo pair CC2703/CC2705. This fragment was cloned in pHM2-Pspac^con^ between BamHI and SalI restriction sites.

The *ytsJ*-GFP translational fusions were made by amplification by PCR of a DNA fragment starting at the +1 promoter of the P2*-ytsJ* mRNA and covering the first 96 AA of the YtsJ CDS using oligo pair CC2815/CC2822. The 3’ UTR of *ytsJ* with its transcriptional terminator was amplified in a PCR reaction using oligo pair CC2823/CC2816. The GFP fragment was amplified using oligo pair CC2821/CC2824. The three overlapping fragments were amplified in a new PCR reaction using oligo pairs CC2815/CC2816. The resulting product, which is a translational fusion of the GFP after the 96th aa of YtsJ CDS, was cloned in pDG148 between the ClaI and HindIII restriction sites (pl912). Since the level of the GFP was too low to be measurable, we place this fusion under the inducible promoter Pspac^con^ of the pDG148-Pspac^con^ plasmid. The fusion was reamplified from pl912 using oligo pair CC2984/CC2816. The transcriptional start site of this transcript is close to the +1 of the P2 promoter (Pspac(P2)-*ytsJ*-GFP). We also amplified a shorter version of this fusion to get a transcriptional start site close to the +1 from P1 promoter (Pspac(P1)-*ytsJ*-GFP) using oligo pair CC2704/CC2816. Both fusions, Pspac(P1)-*ytsJ*-GFP and Pspac(P2)-*ytsJ*-GFP were cloned into the HindIII restriction site of the pDG148-Pspac^con^ plasmid.

### Medium

Strains were grown in MD medium (23), 2xTY or LB medium as indicated, all supplemented with 0.5% malate.

### Sample preparation for RNAseq and pulsed SILAC experiments

Strains CCB1244 and CCB1243 were grown in 50 ml MD modified supplemented with 0.5 % malate and a limiting concentration of lysine (85.5 μM) with shaking. Cultures were grown to OD_600_ = 0.5 and split in two cultures of 20 mL. For the pulse labelling, 1 % arabinose and 1.7 mM medium ^2^H_4_-lysine or heavy ^13^C_8_-lysine was added to CCB1244 and CCB1243, respectively, for 15 min before harvesting the cultures (10 mL for RNAseq analysis and 10 mL for pulsed SILAC analysis). Experiments were performed in triplicate.

### RNAseq analysis

Total RNA was isolated from 10 mL culture (pelleted and frozen) by the glass beads/phenol method described previously (24). RNA samples were treated with RQ DNase Promega (37°C for 20 min) to remove potential contaminating chromosomal DNA. Ribosomal RNA was removed using the RiboZero kit (Illumina), and ribosomal RNA depletion and overall RNA quality was analysed by Bioanalyser (Agilent). cDNA libraries were prepared using the Smarter Stranded RNA-Seq Kit (Clontech) with adapters for multiplexing, according to the manufacturer’s instructions. cDNA concentration and quality were checked by Bioanalyser (Agilent). The 12 samples were normalised to 2 nM, multiplexed and denatured at a concentration of 1 nM using 0.1 N NaOH (5 min at room temperature) before dilution to 10 pM and loading on a HiSeq Rapid SE65. Reads were mapped by Bowtie 2 (25). The analysis was performed using the R software, Bioconductor(26) packages including DESeq2 (27, 28) and the PF2tools package (version 1.5.3) developed at PF2 (Institut Pasteur). Normalization and differential analysis were carried out according to the DESeq2 model and package (version 1.20.0).

### Pulsed-SILAC samples preparation

Proteins were extracted from 10 mL cultures. Cells were lysed twice with a French press, centrifuged to remove cell debris and proteins were precipitated with 5 volumes of ice cold acetone. Protein extracts from the pulse labeling experiments (arabinose-induced RoxS small RNA labeled with heavy ^13^C_8_-lysine and control cultures labeled with medium ^2^H_4_- lysine) were mixed in a 1:1 (w/w) ratio and 25 μg of proteins were loaded on a 12% acrylamide gel prior to separation according to their molecular weight by SDS-PAGE. The gel was fixed and stained with Coomassie Brilliant Blue R250. The gel lanes corresponding to pulsed-SILAC biological triplicates were cut alike into five bands and subjected to a manual in-gel digestion with modified porcine trypsin (Trypsin Gold, Promega). After destaining, bands were subjected to a 30 minutes reduction step at 56°C using 10 mM dithiotreitol in 50 mM ammonium bicarbonate (AMBIC) prior to a 1-hour cysteine alkylation step at room temperature in the dark using 50 mM iodoacetamide in 50 mM AMBIC. After dehydration under vacuum, bands were re-swollen with 250 ng of trypsin in 50 mM AMBIC and proteins were digested overnight at 37°C. Supernatants were kept and peptides present in gel pieces were extracted with 1% (v/v) trifluoroacetic acid. Corresponding supernatants were pooled and dried in a vacuum concentrator. Peptide mixtures were solubilized in 25 μL of solvent A (0.1% (v/v) formic acid in 3% (v/v) acetonitrile) for mass spectrometry analysis.

### Tandem mass spectrometry of SILAC-labeled protein extracts

Mass spectrometry analyses were performed on a Q-Exactive Plus hybrid quadripole-orbitrap mass spectrometer (Thermo Fisher, San José, CA, USA) coupled to an Easy 1000 reverse phase nano-flow LC system (Proxeon) using the Easy nano-electrospray ion source (Thermo Fisher). Each peptide mixture was analyzed in duplicate. Five microliters were loaded onto an Acclaim PepMap™ precolumn (75 μm × 2 cm, 3 μm, 100 Å; Thermo Scientific) equilibrated in solvent A and separated at a constant flow rate of 250 nL/min on a PepMap™ RSLC C18 Easy-Spray column (75 μm × 50 cm, 2 μm, 100 Å; Thermo Scientific) with a 90 min gradient (0 to 20% B solvent (0.1% (v/v) formic acid in acetonitrile) in 70 min and 20 to 37% B solvent in 20 min).

Data acquisition was performed in positive and data-dependent modes. Full scan MS spectra (mass range m/z 400-1800) were acquired in centroïd with a resolution of 70,000 (at m/z 200) and MS/MS spectra were acquired in centroid mode at a resolution of 17,500 (at m/z 200). All other parameters were kept as described in (29)

### Data processing and SILAC quantification

Raw data were processed using the MaxQuant software package (http://www.maxquant.org, version 1.5.6.5) (30). Protein identifications and target decoy searches were performed using the Andromeda search engine and the UniprotKB database restricted to *Bacillus subtilis* taxonomy (release: 01/2019; 4260 entries) in combination with the Maxquant contaminant database (number of contaminants: 245). The mass tolerance in MS and MS/MS was set to 10 ppm and 20 mDa, respectively. Methionine oxidation, protein N-terminal acetylation as well as heavy ^13^C_8_-lysine and medium ^2^H_4_-lysine were taken into consideration as variable modifications whereas cysteine carbamidomethylation was considered as fixed modification. Trypsin was selected as the cutting enzyme and a maximum of 2 missed cleavages were allowed. Proteins were validated if at least 2 unique peptides with a peptide FDR < 0.01 were identified. The setting “Match between runs” was also taken into consideration to increase the number of identified peptides. For quantification, we only used unique peptides with a minimum ratio count ≥ 1. Protein ratios were calculated as the median of all peptide ratios for the protein of interest and a normalisation step was also included in the quantification process.

### Statistical analysis of the pulsed-SILAC dataset

Statistical analysis was done using Perseus software (https://maxquant.org/perseus, version 1.6.0.7) on normalized protein ratios. Proteins belonging to contaminant and decoy databases were filtered. For each biological replicate, the median ratios (M/L and H/L) of the two injected replicates were determined. Then, a student t-test (threshold p-value < 0.05) was applied to detect statistically relevant variations between these two ratios. Moreover, we considered only (H/L)/(M/L) ratios having a fold change >1.4 for up-regulated gene and <0.7 for downregulated gene to validate newly synthetized proteins that are specific to the arabinose-induced RoxS sRNA condition.

All LC-MS raw data files as well as Maxquant and Perseus result files have been deposited to the MassIVE repository with the dataset identifier MSV000091084 and can be freely downloaded at the following address:

https://massive.ucsd.edu/ProteoSAFe/dataset.jsp?accession=MSV000091084.

### IntaRNA prediction

The following parameters have been used for the intaRNA prediction. The output parameters were defined as follows: number of interactions per RNA pair: 5; Suboptimal interaction overlap: can overlap in target; Max absolute energy of an interaction: 0, Max delta energy above mfe of an interaction: 100, no lonely base pairs and no GU helix ends. The seed parameters were: Minimum of basepairs in seed: 7. The folding parameters were: Temperature for energy computation: 37°C; folding window size: 150; max. base-pair distance: 100; folding window size: 150; base-pair distance 100; energy parameter set: Turner model 2004. For each prediction, the target sequence encompasses the proximal 5’ and 3’ end of each mRNA determined in Nicolas *et al*.(15). When the putative target gene is the only gene found to be regulated in an operon, for example ClpQ, 100 nts were added to the 5’ and 3’ ends of the ORF for prediction.

### Gene Ontology analysis

The 83 downregulated genes and the 19 upregulated genes identified by RNAseq and SILAC were used for a Gene ontology analysis on the web application “Comparative Go” (12). A hypergeometric test was used to calculate under-represented or enriched biological functions in these dataset with a p-value <0.05.

### GFP measurement

Strains were grown overnight in LB medium with 0.5% malate and 5 μg/ml phleomycin. The cells were diluted the following day in the same medium. At mid-exponential phase (O.D._600_=0.6-0.8) cells were inoculated at (O.D._600_=0.01) on 96-well microplates and the O.D._600_ and GFP measured for 15 hours. The GFP arbitrary units were corrected with a control strain (CCB1133) with no GFP and an interrupted *amyE* gene, since we observed that the AmyE protein increases fluorescence background during growth.

### Electrophoretic mobility shift assays (EMSA)

For EMSA assays, RoxS was transcribed with T7 RNA polymerase *in vitro* from PCR templates amplified using the oligo pair CC1482/CC1833. The RoxS CCC to GGG mutation in CRR1(RoxS CRR1^/3G^) was transcribed from a PCR template amplified using oligo pair CC2831/1833. The RoxS CCC to GGG mutation in CRR3 (RoxS CRR3^/3G^) was transcribed from PCR templates amplified using oligo pairs CC1482/CC2929 and CC2928/CC1833. The overlapping fragments were then assembled in a new PCR reaction and amplified using CC1482/CC1833.

The *ycsA* and the *ycsA*^m^ RNA were transcribed *in vitro* from PCR templates amplified using the oligo pairs CC2644/CC2642 and CC2987/CC2642 respectively. The *dnaE* RNA was transcribed *in vitro* from a PCR template amplified using the oligo pair CC2645/CC2651. The *dnaE*^m^ template was made in by reamplification of overlapping PCR fragments amplified using oligo pairs CC2645/CC3179 and CC3178/CC2651. The reamplification step was done using oligos CC2645/CC2651.

Before addition to EMSA assays, each RNA was individually heated for 3 min and cooled to room temperature for 10 min. A 15 μL reaction was prepared by mixing 5 pmol of target RNA with increasing concentrations of RoxS (2.5, 5 and 10 pmol) in 1xRNA binding Buffer (10 mM Tris pH 8; 50 mM NaCl; 50 mM KCl, 10 mM MgCl_2_) and incubated at 37°C. After 10 min of incubation, 5 μL of glycerol (stock solution 80%) was added and RNAs were loaded on a 6% non-denaturing polyacrylamide gel (acrylamide:bisacrylamide ratio 37.5:1). Following migration (2h30 at 230V in cold room), RNA was transferred to a Hybond N+ membrane and hybridized with the radiolabelled probe.

### RNA isolation and northern blot

sRNA was isolated from mid-log phase *B. subtilis* cells growing in 2YT, LB or MD medium either by the glassbeads/phenol method described in (24) or by the RNAsnap method described in (31). Typically 5 μg RNA was run on 1% agarose or 5% acrylamide gels and transferred to hybond-N membranes (Cytiva) at in 0.5X TBE Buffer (100V for 4h at 4°C). Hybridization was performed using 5’-labeled oligonucleotides or riboprobe labelled with P^32^-UTP using Ultra-Hyb (Ambion) hybridization buffer at 42°C for a minimum of 4 hours. Membranes were washed twice in 2XSSC 0.1% SDS (once rapidly at room temperature (RT) and once for 10 min at 42°C) and then 3 times for 10 mins in 0.2XSSC 0.1% SDS at RT. Oligonucleotides used are shown in Table S3.

## ACKNOWLEDGMENTS

This work was supported by funds from the CNRS (UMR8261), Université Paris Cité and the French National Research Agency by ANR-16-CE12-0002 (BaRR) and ANR-18-CE12-0025 (CoNoCo). This work was also supported by LABEX DYNAMO (ANR-LABX-011) and EQUIPEX CACSICE (ANR-11-EQPX-0008), notably through funding of the Proteomic Platform of IBPC (PPI).

We thank Jennifer Hermann for the kind gift of the YisK antibody.

## Figure Legends

Supplementary Figure 1: Growth of the control strain (*lysA-, ΔroxS*, Pxsa-Ter) and RoxS over-expressing strain (*lysA-, ΔroxS*, Pxsa-RoxS) in MD medium with or without addition of lysine and arabinose at the mid-exponential phase.

Supplementary Figure 2: Predicted RoxS binding sites in mRNAs encoding enzymes of the TCA cycle or the glycine decarboxylation pathway. The +1 position on mRNA targets and on RoxS correspond to the AUG codon and the transcriptional start site respectively. Binding energies are given underneath the predicted base-pairing.

Supplementary Figure 3: Predicted RoxS (left) or FsrA (right) binding sites in the *lutABC* mRNAs and *pycA* mRNA. Binding energies are given underneath the predicted base-pairing.

Supplementary Figure 4: Predicted RoxS binding sites in the *amhX, clpQ, rpmE* and *yjiC* mRNAs. Binding energies are given underneath the predicted base-pairing.

Supplementary Figure 5: Mapping of the 5’ end of the D2 degradation intermediate protected by RoxS. (**A**.) Primer extension on total mRNA extracted from the WT, *ΔroxS, ΔrnjA, ΔrnjA ΔroxS, Δrny* and *Δrny ΔroxS* strains overexpressing *dnaE* at the *amyE* locus, using a primer specific for *dnaE*. (**B**.) Primer extension on total mRNA extracted from the WT, *ΔroxS, ΔrnjA, ΔrnjA ΔroxS, Δrny* and *Δrny ΔroxS* strains, using a primer specific for *ytsJ*. The RoxS binding site is indicated by a red square. * indicates the 5’ end of D2 intermediate and suspected RNase Y cleavage site.

Supplementary Figure 6: Measurement of half-life of *ytsJ* transcripts in MD + malate (0.5%) medium. Northern blot of total RNA from WT and *ΔroxS* mutant strain isolated at different times after addition of rifampicin, probed for *ytsJ* mRNA. The blot was re-probed for 16S rRNA as a loading control.

Supplementary Figure 7: Potential RoxS binding sites in coding sequences of gene affected by RoxS overexpression. Potential binding site are shown in red and numbered according to the AUG start codon.

Table S1: Strains and plasmids used in this study.

Table S2: Oligonucleotides used in this study.

