## Supplementary material for "New RoxS sRNA targets identified in *B. subtilis* by pulsed SILAC": Table S1

**Strains**

| Name | Genotype | References |
|---|---|---|
| SSB1002 | W168 *trpC2* | |
| CCB441 | *rny::spc* | Durand et *al.* 2012 |
| CCB448 | *rnjA::kan* | Durand et *al.* 2012 |
| CCB485 | *roxS::kan* | Durand et *al.* 2015 |
| CCB558 | *roxS::kan rny::spc* | Durand et *al.* 2015 |
| CCB559 | *roxS::kan rnjA::kan* | Durand et *al.* 2015 |
| CCB1243 | *lysA::ery amyE::pDG1662-Pxsa-RoxS roxS::kan* | This study |
| CCB1244 | *lysA::ery amyE::pDG1662-Pxsa roxS::kan* | This study |
| CCB1314 | *roxS::kan amyE* ::pDG1662-Pxsa pDG148-*ycsA* -GFP | This study |
| CCB1423 | *amyE* ::pDG1662-PxsA pDG148-*ycsA* -GFP | This study |
| CCB1403 | *amyE* :pHM2-Pspac(con)-*dnaE* | This study |
| CCB1404 | *roxS::kan amyE* :pHM2-Pspac(con)-*dnaE* | This study |
| CCB1411 | *rnjA::spc amyE* :pHM2-Pspac(con)-*dnaE* | This study |
| CCB1412 | *roxS::kan rnjA::spc amyE* :pHM2-Pspac(con)-*dnaE* | This study |
| CCB1413 | *rny::spc amyE* :pHM2-Pspac(con)-*dnaE* | This study |
| CCB1414 | *roxS::kan rny::spc amyE* :pHM2-Pspac(con)-*dnaE* | This study |
| CCB1590 | *amyE::* pDG1662-PxsA pDG148-Pspac(P2)-*ytsJ* -GFP | This study |
| CCB1591 | *roxS::kan amyE::* pDG1662-PxsA pDG148-Pspac(P2)-*ytsJ* -GFP | This study |
| CCB1592 | *amyE::* pDG1662-PxsA pDG148-Pspac(P1)-*ytsJ* -GFP | This study |
| CCB1593 | *roxS::kan amyE::* pDG1662-PxsA pDG148-Pspac(P1)-*ytsJ* -GFP | This study |

**Plasmids**

| | Name | Description |
|---|---|---|
| pl805 | pDG1662-Pxsa-RoxS | Arabinose inducible promoter of xsa gene cloned in pDG1662. Allows cloning in +1 (SpeI site) |
| pl801 | pDG1662-Pxsa | Integrative plasmid (*amyE* ) containing RoxS under control of arabinose dependent promoter Pxsa. Allows tight expression from +1. RoxS cloned between SpeI and EcoR1 restriction sites. |
| pl862 | pDG148-*ycsA* -GFP | CC2546/2547 and 2548/572. 2nd PCR amplification with CC2546/572. Cloned into the EcoRI/BamHI restriction sites of pHM2. The *ycsA* -GFP fusion from plasmid 854 was recloned in pDG148; oligos CC2611/2612. Cloned in EcoR1/BamHI |
| pl886 | Pspac(con)-*dnaE* | From internal P2 promoter to end of *dnaE* . coordinates 2992494-2991279. Amplified by PCR (CC2703/2744). Cloned into BamHI/SalI |
| pl934 | pDG148-Pspac(P2)-*ytsJ* -GFP | Fusion *ytsJ* -GFP. long 5'UTR (P2). GFP fusion at K(96). cloned into the HindIII restriction site. Under control of the Pspac. |
| pl935 | pDG148-Pspac(P1)-*ytsJ* -GFP | Fusion *ytsJ* -GFP. short 5'UTR (P1). GFP fusion at K(96) cloned into the HindIII restriction site. Under control of the Pspac. |
