## Supplementary material for "New RoxS sRNA targets identified in *B. subtilis* by pulsed SILAC": Table S2

**Oligos**

| Name | Sequence | Description |
|---|---|---|
| | | **for Northern blot** |
| CC2166 | CAAAGGGTTTTAGTTCCAGCAGC | Oligo fwd. for synthesis of *citZ* riboprobe |
| CC2167 | GCTCTAATACGACTCACTATAGGGATACCCCACATATGTAAGGGTATCATC | Oligo Rev. with T7 promoter for synthesis of *citZ* riboprobe |
| CC2577 | GCTCTAATACGACTCACTATAGCTCTGTCCTGGCACCTGAAAG | Oligo Rev. with T7 promoter for synthesis of *gcvT* riboprobe |
| CC2578 | CATAACAGCATGAAAATATGAGCG | Oligo fwd. for synthesis of *gcvT* riboprobe |
| CC3062 | GCTCTAATACGACTCACTATAGGCAAGCTGTCCGAGATAGAAATC | Oligo Rev. with T7 promoter for synthesis of *lutB* riboprobe |
| CC3063 | GAGCGGGTATCACAGGGGATTG | Oligo fwd. for synthesis of *lutB* riboprobe |
| CC2533 | GCTCTAATACGACTCACTATAGGGCATTCTGGATGGCGATTTCATC | Oligo Rev. with T7 promoter for synthesis of *ycsA* riboprobe |
| CC2534 | CTTGGAGCACGGCAAAATGATGC | Oligo fwd. for synthesis of *ycsA* riboprobe |
| CC2826 | GCTCTAATACGACTCACTATAGGGCGCCGCTTCAGGGCCGATGTTTCC | Oligo Rev. with T7 promoter for synthesis of *ytsJ* riboprobe |
| CC2090 | GGAGAATGTCATTAAGAGAAGAAG | Oligo fwd. for synthesis of *ytsJ* riboprobe |
| | | **for primer extension** |
| CC2651 | GCGCGGAGGCTGTATCTGACAGGCGCGGAGGCTGTATCTGACAG | Oligo for mapping of of the P2 promoter of *ytsJ* in *dnaE* |
| CC2109 | GCCGATGTTTCCAAGGCCAAGCAC | Oligo for mapping of of the P1 promoter of *ytsJ* in *dnaE* |
| | | **for strain construction** |
| CC2250 | GACGAAGGATCCCATATTTATAAATACATACGTAC | Oligo fwd. to clone the xsa (abf2) promoter inducible with arabinose in pDG1662. |
| CC2251 | GACGAAAAGCTTGTAAGCGCTTTTACTAGTATTATATTATATATGTTC | Oligo rev. to clone the xsa (abf2) promoter inducible with arabinose in pDG1662. |
| CC2285 | GTCGTGACTAGTGTGAAATTGATCACAAACAAAC | Oligo fwd. to clone RoxS under the inductible arabinose promoter (PxsA) in pDG1662-Pxsa (cloning site SpeI). |
| CC2286 | GTCGTGGAATTCCCAAACGTACGTTCCATTTTTAGCAC | Oligo rev. to clone RoxS under the inductible arabinose promoter (PxsA) in pDG1662-Pxsa (cloning site SpeI). |
| CC2546 | ACATGAGAATTCTTAATCAATAACCGACCACCCG | Oligo fwd. to amplify *ycsA* in order to construct the *ycsA*-GFP fusion (cloning site EcorI) |
| CC2547 | GACAACTCCAGTGAAAAGTTCTTCTCCTTTACTGAGCTTTCTTAAGCGCGAACAG | Oligo rev. to amplify *ycsA* in order to construct the *ycsA*-GFP fusion |
| CC2548 | CTGTTCGCGCTTAAGAAAGCTCAGTAAAGGAGAAGAACTTTTCACTGGAGTTGTC | Oligo fwd. to amplify gfp. Complementary to CC2547 |
| CC572 | AAGCAAAGGGATCCCCTGTCAGGAAGCCCATGCTC | Oligo rev. In pHM2 to amplify GFP from pHM2-5'hbs-GFP plamid (cloning site BamHI) |
| CC2611 | ACATGAATCGATTTAATCAATAACCGACCACCCG | Oligo fwd. to amplify *ycsA*-GFP construct from pHM2-*ycsA*-GFP to clone it in pDG148 (cloning site EcorI) |
| CC2612 | AACGAAGGTCGACCCTGTCAGGAAGCCCATGCTC | Oligo rev. to amplify *ycsA*-GFP construct from pHM2-*ycsA*-GFP to clone it in pDG148 (cloning site PvuII) |
| CC2703 | ACACAGGATCCACACTGCAAATGAGGTCTACG | Oligo fwd. to clone the dnaE fragment into pHM2-Pspac© (cloning site BamHI) |
| CC2705 | ATATAGGATCCGACAAGGTTGACAGCGCATTC | Oligo rev. to clone the dnaE fragment into pHM2-Pspac© (cloning site SalI) |
| CC2815 | TATAATCGATGGGCGGTCAGCAAAAAGAAAAAAG | Oligo Fwd. to construct the *ytsJ*-GFP fusion (from P2 promoter) |
| CC2822 | CTCCAGTGAAAAGTTCTTCTCCTTTACTTTTGAAAAGAACTGCTTTCCCTTCC | Oligo rev. overlap GFP-*ytsJ* to construct the *ytsJ*-GFP fusion at the 96 aa of *ytsJ* (from P2 promoter). |
| CC2821 | GGAAGGGAAAGCAGTTCTTTTCAAAAGTAAAGGAGAAGAACTTTTCACTGGAG | Oligo fwd. overlap GFP-*ytsJ* to construct the *ytsJ*-GFP fusion at the 96 aa of *ytsJ* (from P2 promoter). |
| CC2824 | GCGCTGCCATTTTTGGAATGAATTAAAATTATTTGTATAGTTCATCCATGCCATGT | Oligo Rev. overlap ytsJ-GFP pour clonage des fusions ytsJ-GFP. |
| CC2823 | ACATGGCATGGATGAACTATACAAATAATTTTAATTCATTCCAAAAATGGCAGCGC | Oligo Fwd. overlap GFP-*ytsJ* pour clonage des fusions ytsJ-GFP. (3'UTR ytsJ) |
| CC2816 | TATAAAGCTTATTCGATGACGTGCTCGGAAAGG | Oligo Rev. to clone the *ytsJ*-GFP fusion into pDG148. (3'UTR ytsJ) |
| CC2984 | ATATAAAGCTTACACTGCAAATGAGGTCTACG | Oligo fwd. to amplify the *ytsJ*-GFP fusion from pDG148-P2-*ytsJ*-GFP and cloned it into the pDG148-Pspac(c). Fusion Pspac(P2)-*ytsJ*-GFP |
| CC2704 | ACACAGGATCCAATTATGTGATCCTGTTTAAATATTCAGATAG | Oligo fwd. to amplify the *ytsJ*-GFP fusion from pDG148-P2-*ytsJ*-GFP and cloned it into the pDG148-Pspac(c). Fusion Pspac(P1)-*ytsJ*-GFP |
| | | **for EMSA** |
| CC2644 | TAATACGACTCACTATAGGCGCTTACGATAACAGGGGAAGG | Oligo fwd. with T7 promoter for synthesis of *ycsA mRNA* |
| CC2987 | TAATACGACTCACTATAGGCGCTTACGATAACAGCCCAAGGAGAATGACGATG | Oligo fwd. with T7 promoter for synthesis of *ycsAm mRNA* |
| CC2642 | CCGCTGTATGAAGCACTTTCTCAGC | Oligo rev. for synthesis of *ycsA and ycsAm* (3G->3C) mRNA |
| CC2645 | GCTCTAATACGACTCACTATAGGACACTGCAAATGAGGTCTAC | Oligo fwd. with T7 promoter for synthesis of *dnaE mRNA* |
| CC2651 | GCGCGGAGGCTGTATCTGACAGGCGCGGAGGCTGTATCTGACAG | Oligo rev. for synthesis of *dnaE and dnaEm* mRNA |
| CC3178 | CAATATCTTTACGAAGCAAAGCCCAGCGGGATTCGTATCC | Oligo fwd. for synthesis of *dnaE mRNA* (3G->3C) |
| CC3179 | GGATACGAATCCCGCTGGGCTTTGCTTCGTAAAGATATTG | Oligo rev. for synthesis of *dnaE mRNA* (3G->3C) |
| CC1482 | GCTCTAATACGACTCACTATAGGTGAAATTGATCACAAACAAACATTAC | Oligo fwd. with T7 promoter for synthesis of RoxS sRNA |
| CC1833 | AAAGAAACCGCGCCGGGATAAG | Oligo rev. for synthesis of RoxS sRNA |
| CC2831 | GCTCTAATACGACTCACTATAGTGAAATTGATCACAAACAAACATTACGGGTTTGTTT GACCGTG | Oligo fwd. with T7 promoter for synthesis of RoxS CRR1 sRNA (3C->3G) |
| CC2928 | GTGAAAAAATTTCTCCCATGGGCTTTGTTGTCGTTAAG | Oligo fwd. for synthesis of RoxS CRR3 sRNA (3C->3G) |
| CC2929 | CTTAACGACAACAAAGCCCATGGGAGAAATTTTTCAC | Oligo rev. for synthesis of RoxS CRR3 sRNA (3C->3G) |
